# Distinct Control of histone H1 expression within the Histone Locus body by CRAMP1

**DOI:** 10.1101/2025.01.07.631602

**Authors:** Justin Bodner, Pranathi Vadlamani, Kathryn A. Helmin, Qianli Liu, Marc L. Mendillo, Benjamin D. Singer, Shashank Srivastava, Daniel R. Foltz

## Abstract

Proper histone gene expression is critical to cell viability and maintaining genomic integrity. Multiple histone genes organized into three genomic loci encode for replication coupled core and linker histones. Histone gene expression and transcript processing is orchestrated in the histone locus body (HLB) within the nucleus. We identified human CRAMP1 as a selective regulator of linker histone H1 expression. CRAMP1 is recruited to the HLB in RPE1^hTERT^ cells. Affinity purification shows that CRAMP1 physically associates the HLB component GON4L (a.k.a. YARP). We show that the PAH domains of GON4L interact with CRAMP1. CRAMP1 disruption results in a loss of histone H1 expression and a reduction in H1 protein. CRAMP1 occupies the unmethylated promoters of the replication coupled linker histone genes that reside within the histone locus body, and the replication independent histone H1 loci, which reside in a region of the genome without other histone genes. Together these data identify CRAMP1 as a novel and selective regulator of histone H1 gene expression.

## Introduction

The human genome includes over 90 histone genes that encode the core (H2A, H2B, H3 and H4) and H1 linker histones. Most human genes encoding the replication coupled histones are arranged into 4 clusters spread across chromosomes 1,6, and 12 ^1–4^. The locus on human chromosome 6 is the largest of the clusters, containing approximately 70% of all histone genes and 66% of linker histone genes. The multimerization of the histone genes is conserved from S. *cerevisiae* to humans, but the arrayed format of histone genes is not. In *Drosophila*, histone genes pairs are organized in a head-to-head arrangement and exist as ∼100 tandem repeat units, each of which includes an H3/H4 and H2A/H2B pair and a single H1 gene ^5^. The divergent head-to-head arrangement of histones observed in model organisms is poorly conserved in humans. However, the organization of the histone genes into these 4 clusters is syntenic across mammals^1^.

Histone gene expression is controlled at both transcriptional and post transcriptional levels^6^. In higher eukaryotes, replication coupled histone genes lack introns, and most of the transcripts do not undergo polyadenylation. Instead, the 3’UTR regions form well conserved, stem-loop structures. The stem loop binds to SLBP, and in combination with the U7 snRNP, positions the CPSF73 exonuclease to cleave the 3’ end of the nascent transcript. When histone H2A-H2B gene expression is increased by just 2-fold, the overall RNA level of the histones is maintained at an unchanged level by enhanced RNA turnover^7^. Misregulation of histone gene expression leads to defects in cellular fitness and growth ^8–11^. Overexpression of individuals histones, or subsets of the nucleosome in S.*cerevisiae* leads to a reduction of histone gene expression^12^.

The histone locus body (HLB) is a cytologically defined nuclear body that is the site of histone mRNA biogenesis^13–16^. HLBs are membraneless organelles and liquid-liquid phase separation is proposed to play a role in their formation^17,18^. In D*rosophila*, sequences within the histone H3/H4 gene regions of the tandem repeats are sufficient to establish a histone locus body^19^. Expression of the core histone genes requires the protein NPAT (Nuclear Protein, Ataxia-Telangiectasia/ Nuclear Protein, Mxc in *Drosophila, SPT21 in S.cerevisiae*) ^20,21^. NPAT is a central regulator of HLB formation and core histone expression ^22–24^. The cell cycle restriction of histone gene expression, coupled with DNA replication is controlled thorough CDK phosphorylation of NPAT ^25,26^. Human GON4L (a.k.a. YARP, YY1 Associated associated-related protein) is the homolog of the *Drosophila* Mute gene. GON4L and Mute are recruited to the histone locus bodies in HeLa cells and *Drosophila*, respectively^22^. Mute in Drosophila appears to suppress histone gene expression ^27^; however, it has not been tested if GON4L plays a similar repressive role to the core histone in human cells, and its role in histone H1 regulation has not been tested in either system. GON4L is known to bind the Ying-Yang1 (YY1) transcription factor, and YY1 may play a cell type specific role in histone transcription ^28–32^.

In contrast to the core histones, all histone H1 genes encode for distinct linker histone proteins^33^. Histones H1-1, H1-2, H1-3, H1-4, H1-5 are expressed primarily during S-phase to support DNA replication, and these genes are located with the histone loci on chromosome 6 and occupy the HLB. Germline specific subtypes include H1-6, H1-7 and H1-8. Histones H1-0 and H1-10 are expressed throughout the cell cycle, independent of DNA replication. The replication coupled H1 genes share the highest degree of identity (∼76% – 86%), whereas the replication independent variants share only 30-43% identity with any other homolog. Like the replication-independent histone H3 and H2A variants, H1-0 and H1-10 genes lie outside of the histone clusters. All H1 proteins contain a globular domain with variable N and C termini. Histone H1 interacts with the linker DNA at the entry-exit site at or near the dyad axis of the nucleosome to form a structure called the chromatosome. Many roles for H1 have been identified including chromatin condensation and 3D genome organization ^34^ ^35^.

Core histones use a system of chaperone proteins to facilitate the assembly of nucleosomes^36^. Histone H3 variant-specific chaperones determine the timing and location of variant nucleosome assembly. Whether a unique histone chaperone network exists for histone H1 is not clear, though NASP and NPM1 have been proposed to bind to H1 ^37–39^, and NASP and NPM1 can contribute to H1 assembly in vitro ^37,40^.

Here we identified human CRAMP1 as a regulator of histone H1 gene transcription and a novel member of the human HLB. Affinity purification of CRAMP1 identified the GON4L HLB component as a binding partner of CRAMP1. The interaction between CRAMP1 is mediated by the PAH domains of GON4L. We show that CRAMP1 selectively regulates histone H1 genes, without affecting core histone gene expression. ChIP-seq experiments demonstrate that H1 binds almost exclusively to the promoters of histone H1 genes. Both replication-coupled genes and replication independent histone H1 genes are bound by CRAMP1. Together, these data suggest that the core histone and H1 genes are differentially regulated through CRAMP1, and that diverse levels of regulation occur within the histone locus body.

## Results

### Genome-wide cell fitness analysis identifies CRAMP1 as an H1 regulator

We used a human genetic approach to identify human specific pathways related to histone H1 function by analyzing large-scale cell fitness screens in response to whole-genome CRIPSR/Cas9 drop out screening across >1000 human cell lines from DepMap^41,42,43,44^. Identification of gene knockouts that demonstrate a correlated degree of fitness across a large number of cancer cell lines has been used by us an others to identify physical protein complexes and related regulatory pathways that influence stress response and define complexes involved in chromatin remodeling^45,46^ (Fig.1A).

**Figure 1.**
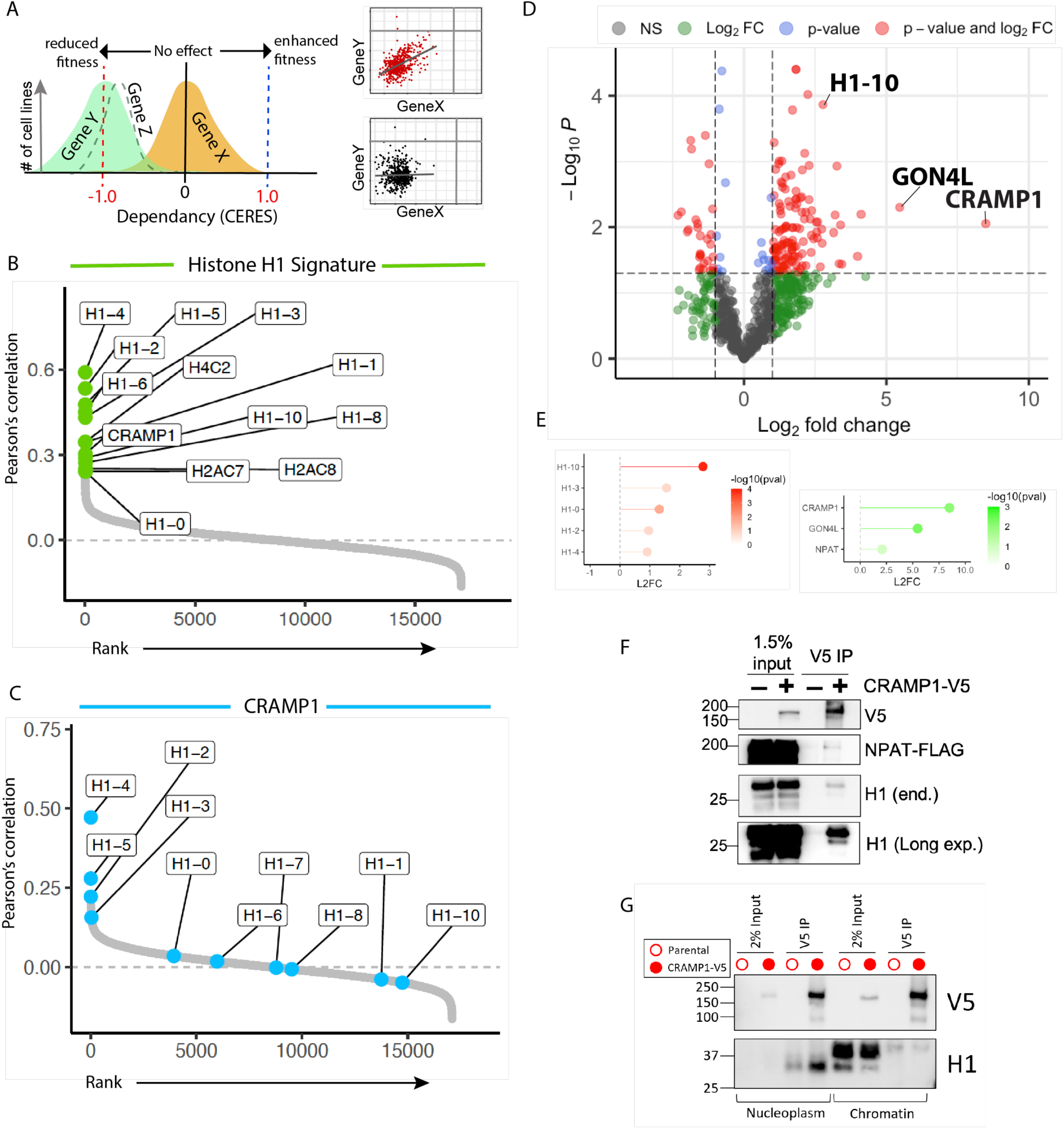
Association of human CRAMP1 and histone H1. A) Schematic showing the use of fitness correlations from whole genome knockouts across multiple cell lines for identification of functionally related genes. B) Rank order of gene correlations with a combined histone H1 fitness signature across the entire genome. CRAMP1 is the highest correlated non-histone protein. C) Rank order of fitness effects of genes correlated with CRAMP1 induced cell fitness. D) HEK293T cells stably expressing CRAMP1-V5 were used to conduct affinity purification of CRAMP1 and proteins were identified by mass spectrometry. n=3. E) Results from affinity purification of individual histone H1 genes and regulators of histone gene expression. F) Co-immunoprecipitation from HEK293T cells stably expressing CRAMP1-V5 and transiently expressing NPAT-FLAG confirming the association of CRAMP1 NPAT and histone H1. G) Fractionation for nucleoplasm and chromatin showing the association of CRAMP1 and H1 in the soluble nucleoplasmic fraction.

To identify genes that are functionally related to histone H1 gene function, we created an H1 gene fitness signature that averaged the fitness effects of all H1 genes across cell lines. CRAMP1 showed the highest correlation with the H1 gene signature of any non-histone gene (P=0.303)^47^ (Fig.1B). CRAMP1 was similarly ranked (P=0.388) when we restricted the gene signature to replication coupled histone H1 genes. Likewise, the replication coupled histone H1 genes (H1-2, H1-4 and H1-5) showed the highest correlation with CRAMP1 (Fig.1C). In addition to the H1 genes, CRAMP1 also showed a high degree of correlation with the PRC1 components, EED, SUZ12, and EZH2. Interestingly, *Drosophila* mutants of the CRAMP1 homolog, Cramped, phenocopy Polycomb mutants^47^.

Affinity purification followed by mass spectrometry (AP-MS) was conducted from an HEK293T cell line stably expressing C-terminally V5-tagged CRAMP1 to identify the proteins associated with CRAMP1 (Fig. 1D). This approach identified 75 enriched proteins in CRAMP1-V5 purification that met the cutoff criteria (Pval < 0.05 and Log2 fold > 1.5) as compared to V5 purification from parental counterpart (Fig.1, red dots). The affinity purification showed CRAMP1 associates with several histone H1 variants. The H1-10 and H1-0 were most abundantly purified, however, histone H1-2, H1-3 and H1-4 were also purified with CRAMP1 (Fig.1D), albeit below the stringency cutoffs assigned (Fig.1E). The co-purification of CRAMP1 with Histone H1 is consistent with the strong fitness correlation between CRAMP1 and the histone H1 genes. In contrast, core histones identified by AP-MS showed no enrichment with CRAMP1-V5 purification.

GON4L (a.k.a YARP) was the most highly purified protein (Log_2_ fold change = 5.5) associated with CRAMP1. GON4L was previously shown to interact with NPAT directly ^22,24^. Although NPAT was also found in our affinity purification-mass spectrometry dataset, it did not meet our cutoff criteria. Nonetheless, we demonstrated that NPAT is co-immunoprecipitated with CRAMP1 when co-expressed in HEK293T cells (Fig.1F), suggesting that this interaction is valid, albeit potentially indirect. We did not identify GON4L in our fitness screening. These incongruencies can occur when genes have multiple phenotypic outcomes that impinge on cellular fitness, therefore, we expect that GON4L will have additional roles in the cell.

Subcellular fractionation was conducted followed by co-immunoprecipitation from HEK293T^CRAMP1-V5^ cells to determine whether CRAMP1 was bound to H1 in the nucleoplasm or was part of the chromatosome, which is histone H1 bound to the nucleosome. CRAMP1 was immunoprecipitated from both the chromatin and soluble nucleoplasmic fractions. However, histone H1 was bound to CRAMP1 predominantly in the nucleoplasmic fraction, suggesting that CRAMP1 is not a structural component of the chromatosome complex.

### Interaction between CRAMP1 and GON4L

CRAMP1 has a well conserved domain structure from *Drosophila* to human. Sequence similarity identified a conserved SANT domain within at the N-terminus of CRAMP1 (a.a. 168-224) (**Fig. 2A**). SANT domains are common protein-protein interaction domains and are present in other proteins of the HLB, including GON4L and FLASH ^22,24^, and can serve as chromatin interaction and remodeling motifs ^48,49^. Based on Alphafold3.0 prediction, the structured region extends beyond the annotated SANT domain to include an additional 77 amino acids.

**Figure 2.**
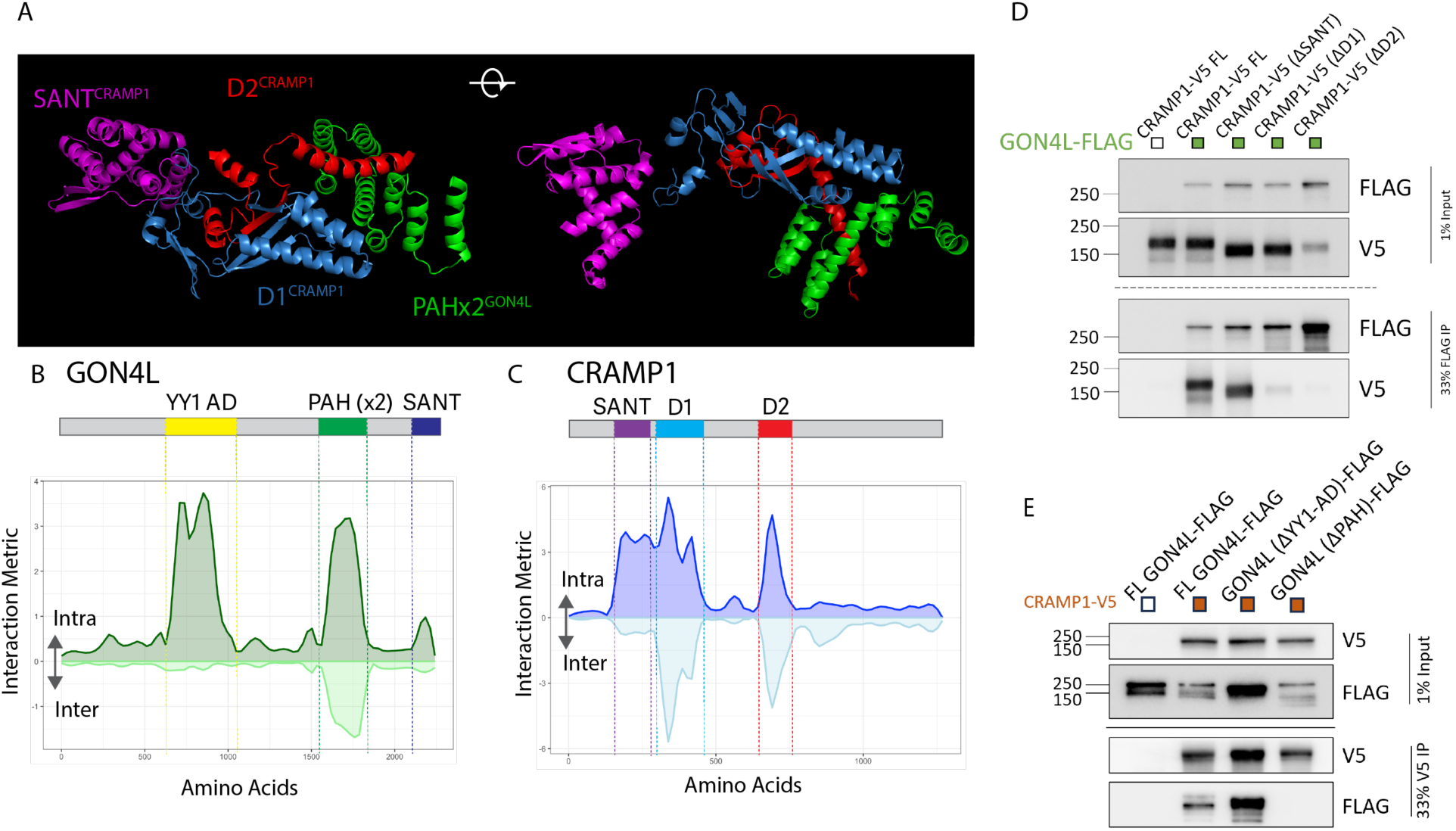
Interaction between CRAMP1 and GON4L. A) AlphaFold 3.0 predicted structure of CRAMP1. Only regions within established or newly defined domains are shown. Sequence predicted and newly identified domains are color coded. B,C) Interaction graphs show the domain structure and predicated amino acids of interaction based on the interaction metric derived from the expected position error. Positive values indicate high confidence structure within the protein and negative values indicate structure between GON4L and CRAMP1. D) Anti-FLAG (GON4L) co-IP from HEK293T cells transiently expressing full length GON4L and indicated CRAMP1 deletion mutants. F) Co-IP of full length CRAMP1-V5 using anti-V5 antibody from HEK293T cells transiently expressing full-length CRAMP-1 and GON4L deletion mutants.

Structural prediction also identified two well-structured regions following the SANT domain in the N-terminal half of the protein (termed D1 (a.a. 315-429) and D2 (a.a. 663-741)) that were not part of a previously recognized motif (Fig 2A,C). Domains D1 and D2 are highly conserved within CRAMP1 across multiple species (Fig.SX). These two domains are separated by a largely unstructured region of 234 amino acids. However, the two regions are predicted to form a common structure (Fig. 2A). Beta-strands from both D1 and D2 contribute to a mixed parallel/anti-parallel beta sheet.

GON4L contains a YY1 binding domain (a.a. 600-1339) and two paired amphipathic helix (PAH) repeats, PAH1 (a.a. 1624-1696) and PAH2 (a.a.1706-1777) (Fig. 2A,B). Modeling demonstrated that two PAH domains in the C-terminus of GON4L may interact with CRAMP1. The 4-helix PAH2 domain of GON4L interacts with the last alpha helix of the D2 domain of CRAMP1 (Fig.2A). The YY1 binding domain of GON4L did not show an interaction interface between GON4L and CRAMP1.

To determine whether the PAH domains of GON4L are required for binding to CRAMP1 we developed FLAG-tagged GON4L deletion mutants lacking amino acids 1585 – 1831, spanning both PAH1 and PAH2, or lacking a.a. 600-1339 that eliminates the YY1 binding domain. Wild-type or CRAMP1 deletion mutants were co-expressed in HEK293T cells in combination with full-length FLAG-tagged GON4L. Full-length CRAMP1 and the deletion mutant lacking the SANT domain efficiently co-precipitated GON4L (Fig. 2D). However, deletion mutants of CRAMP1 that eliminate the D1 (deletion of a.a. 315-439) or D2 domains (deletion of a.a. 663-741) of CRAMP1 abrogated the interaction with GON4L. Likewise, full-length GON4L and the deletion mutant lacking the YY1-binding domain efficiently co-precipitated CRAMP1 (Fig. 2E). In contrast, deletion of the PAH repeats eliminated the interaction between CRAMP1 and GON4L, demonstrating that the PAH domain is required of CRAMP1 to bind GON4L.

### CRAMP1 localization to Histone Locus Bodies (HLB)

To determine the subcellular localization of CRAMP1 in human cells, we created a stable cell line expressing V5-CRAMP1 in RPE1^hTERT^ cells. We observed clear co-localization of CRAMP-V5 with NPAT positive foci (Fig.3A), demonstrating that CRAMP1 is a component of the histone locus body, similar to the Drosophila homolog, cramped (REF). The Cajal Body (CB) and HLB overlap in RPE1-hTERT cells as indicated by Coilin and FLASH staining (Figure SX), similar to many cell types. Consistent with the overlap of the CB and HLB, CRAMP1 and Coilin also co-localized (Fig.3B). We examined the localization of NPAT and CRAMP1 using Airyscan superresolution imaging (Zeiss). NPAT and CRAMP1 occupy distinct and partially overlapping domains within the HLB (Fig.3C). We observed NPAT forms a shell around the HLB and CRAMP1 appears to be largely contained within the core region, similar to what has been observed previously for NPAT localization^50^.

**Figure 3.**
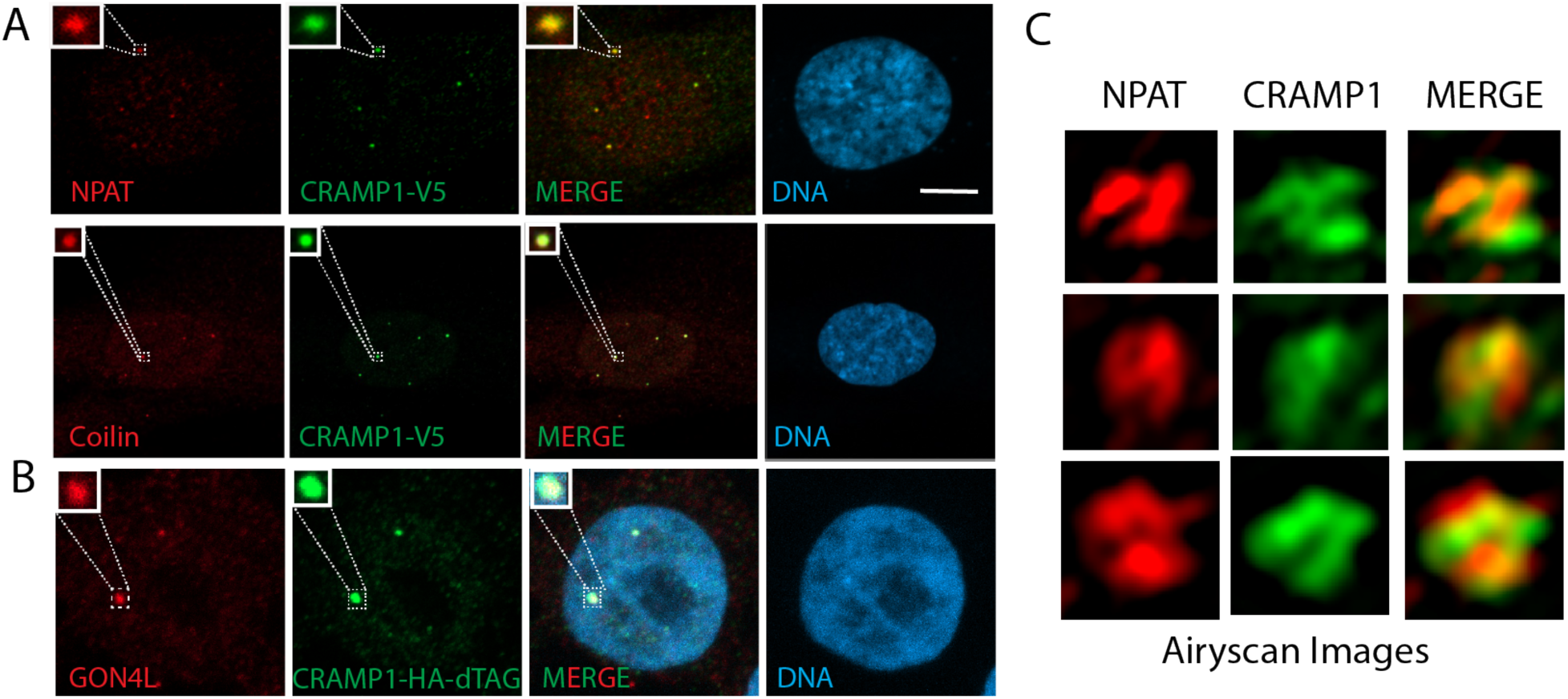
Localization of CRAMP1 to the histone locus body (HLB). A) Immunocytochemistry of RPE1-hTERT cells stably expressing CRAMP1-V5 co-stained with V5, NPAT or Coilin antibodies. CRAMP1-V5 colocalized to the HLB along with NPAT, and with Coilin. Cajal bodies and HLB co-localize in RPE1 cells. B) CRAMP1 was tagged at the endogenous locus in HEK293T cells with a dual HA-dTAG. Endogenous CRAMP1 and GON4L were colocalized at histone locus bodies by immunocytochemistry using antibodies against HA and GON4L. Scale-bar = 5 microns. C) Three high resolution Airyscan images of HLB stained with NPAT and CRAMP1-V5 showing distinct localization of NPAT and CRAMP1.

### CRAMP1 occupies the promoters of H1 genes

Fractionation experiments showed that CRAMP1 was present in both the soluble nucleoplasm and chromatin fraction (Fig 1G), immunofluorescence shows a predominant localization to the HLB. To determine the genomic location of CRAMP1 within the HLB, ChIP-seq was conducted using the RPE1^hTERT^ CRAMP1-V5 expressing cells that show discrete localization of CRAMP1 to the HLB (Fig. 3A). ChIP-seq using an V5 antibody identified 32 peaks common to all three replicates (Fig.4A). The peaks occupied primarily protomer regions and were enriched for genes related to nucleosome structure (Fig.4B,C).

**Figure 4.**
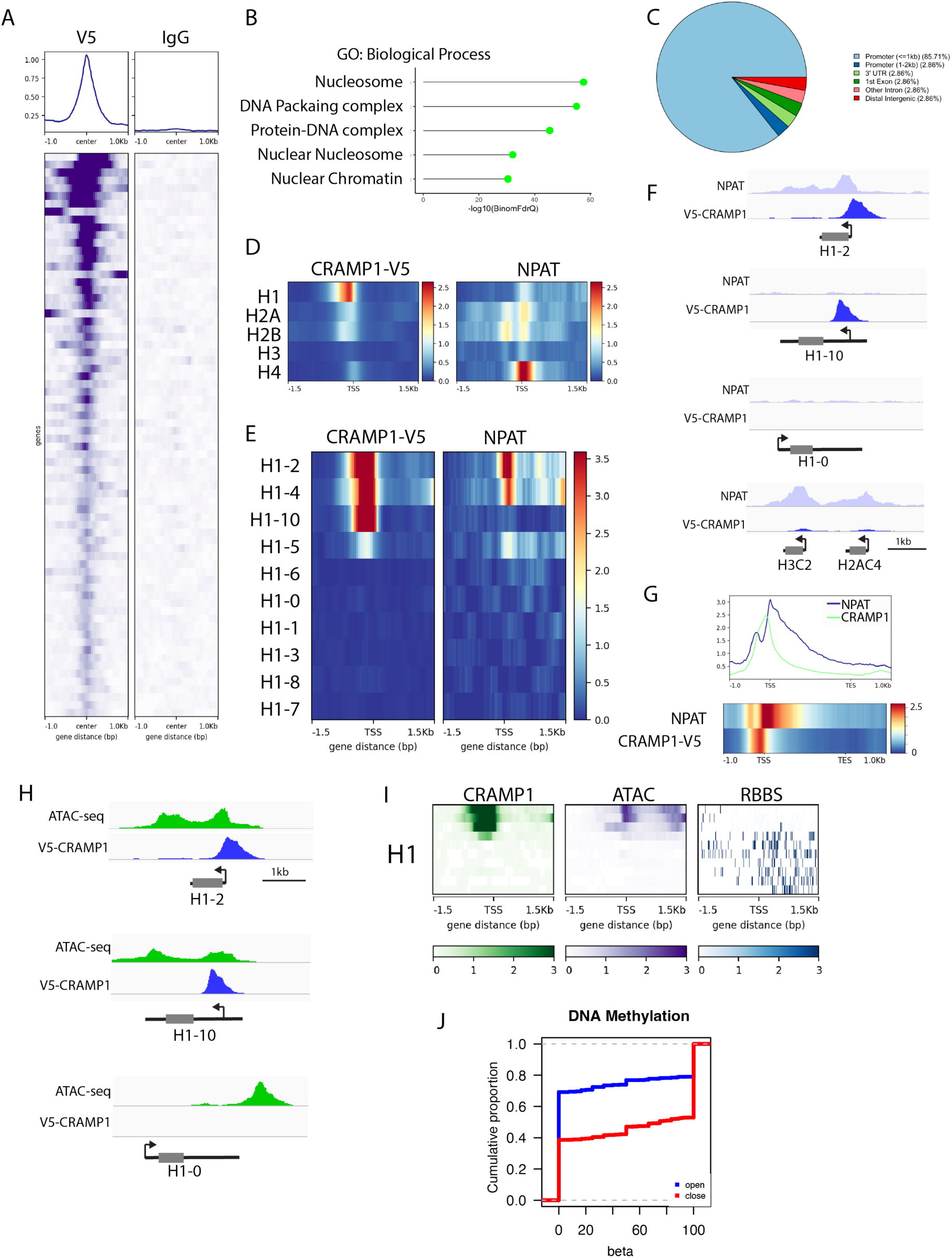
H1 promotors are bound by CRAMP1. A) ChIP-seq was conducted from RPE1^hTERT^ cells stably expressing CRAMP1-V5. Heatmaps display the ChIP-seq signal average of two biological replicates at consensus peaks. B) GO analysis of genes within 1kb of the CRAMP1-consensus binding sites using GREAT identified^58^ enrichment for histone genes. C) Binding sites of CRAMP1 were located primarily within promoter regions. D) Combined heatmaps of all genes within the classes of core (H2A, H2B, H3 and H4) and linker promoters show a preference of CRAMP1 for Histone H1 promoters. E) Heatmaps of V5-ChIP-seq peak intensities at individual histone H1 genes. F) Track examples of genes showing the accumulation of CRAMP-1 from V5 ChIP-seq and NPAT ChIP-seq ^51^. G) Distribution of NPAT and CRAMP1 V5 ChIP-seq signal across the gene. H) Track examples showing the recruitment of CRAMP1-V5 to histone H1 genes that have open chromatin. I) CRAMP1-V5 ChIPseq, ATAC-seq and CpG methylation signals were mapped to the transcriptional start sites of all H1 histone genes. J) Cumulative distribution function of methylation of CpG sites at all histone promoters compared between sites that are categorized as open or closed based on ATAC-seq signal. A leftward shift indicates relative hypomethylation.

We examined the localization of V5-CRAMP1 specifically at the histone genes and compared the CRAMP1 occupancy distribution to a pre-existing NPAT ChIP-seq dataset from RPE1^hTERT^ cells ^51^(Fig.4D). CRAMP1-V5 accumulated to its highest levels at the promoters of the linker H1 genes, with more modest recruitment to H2A, H2B and H4 promoters. In contrast, NPAT, which is expected to regulate the core histones, accumulates most highly at the H4 promoter, and to a lesser degree to H2A, H2B and H1. The biphasic distribution of NPAT at the promoters of H2A and H2B genes reflects the head-to-head arrangement of many H2A and H2B gene pairs within the human genome. Surprisingly, NPAT showed little accumulation at the H3 promoters, suggesting that the influence of NPAT on H3 regulation may be through coordinated control of H3 and H4. CRAMP1 accumulates at the promoter of the replication coupled H1-2 and H-4 genes, as well as the replication independent H1-10 gene. While NPAT is weakly recruited to the H1-2 and H1-4 gene promoters, it is absent from the CRAMP1 accumulated in a discrete region of the promoter, and NPAT is distributed downstream of CRAMP1 and extends into the histone gene body (Fig. 4G).

CRAMP1 is highly recruited to a subset of histone H1 genes, specifically H1-10, H1-2 and H1-4, in RPE1^hTERT^ cells. This is consistent with highly cell type and tissue dependent expression of histone H1 genes. To determine whether CRAMP1 is selectively present at the promoters of active histone H1 genes, we conducted ATAC-seq to identify the open/closed chromatin state at the histone promoters. The ATAC-seq analysis shows a correlation between open-chromatin states at the histone H1 genes and the accumulation of CRAMP1 (Fig. 5H,I), suggesting that CRAMP1 is recruited to accessible histone gene promoters. To define the mechanism by which cells select the histone H1 genes to be expressed, we examined DNA methylation at the promoters of the H1 genes. We observe that DNA methylation negatively correlated with open chromatin and the presence of CRAMP1 at histone gene promoters in general (Fig. 5J). This finding suggests that DNA methylation may be a factor that regulates cell-type specific histone gene expression.

**Figure 5.**
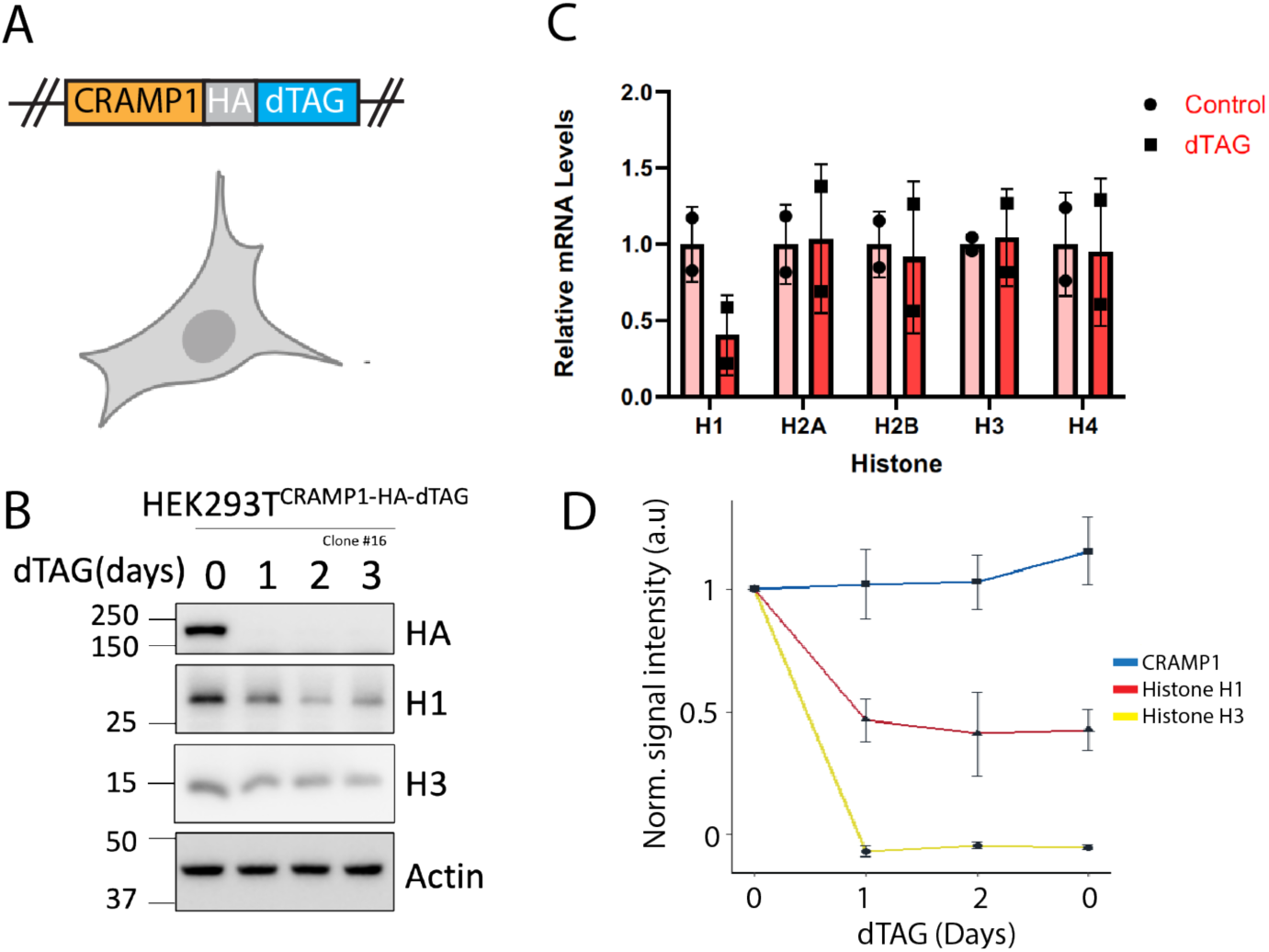
CRAMP1 is required for Histone H1 gene expression. A) A dual HA-dTAG was incorporated at the endogenous locus of the CRAMP1 gene in HEK293T cells. B) Immunoblot showing the protein levels HA-tagged endogenous CRAMP1, Histone H1 and Histone H3. C) HEK293T^CRAMP1-HA-dTAG/CRAMP1-HA-dTAG^ cells were treated DMSO or dTAG13 48 hr followed by RT-PCR for a panel of core and linker histone genes known to be expressed in HEK293T cells. H2A(H2AFY), H2B (H2BC12), H3.1 (H3C6), H4 (H4C9) and H1 (H1-2). Suppression of CRAMP1 selectively reduced histone H1-2 gene expression. n=3. D) Quantitation of protein levels from panel B. n=3.

### Effects of CRAMP1 on histone H1 expression

Since CRAMP1 localizes to the HLB and is associated with two known histone transcriptional regulators, NPAT and GON4L, we sought to determine if CRAMP1 alters histone gene expression. A degron tag (dTAG) was incorporated into the endogenous CRAMP1 locus in HEK293T cells (Fig. 5A). The addition of dTAG13 resulted in the degradation of CRAMP1 to below detectable levels within 1 day of treatment (Fig. 5B). We conducted rtPCR for a subset of histone genes that included one histone from each class of the replication-coupled histone genes located in cluster 1 on chromosome 6 (Figure 5C). Of all the histone genes, only histone H1 showed a reduction in mRNA expression when CRAMP1 was degraded, consistent with the strong fitness correlation between CRAMP1 and histone H1. Coincident with the degradation of CRAMP1, the proteins levels of histone H1 are dramatically reduced (Fig.5B,D), consistent with the function of CRAMP1 in suppressing Histone H1 gene expression.

## Discussion

Here we describe human CRAMP1 as a regulator of histone H1 gene expression. The pattern of cellular fitness across cell types in response to loss of histone H1 is highly correlated with CRAMP1 knockout, demonstrating a functional relationship between Histone H1 and CRAMP1. Consistent with this genetic interaction, CRAMP1 both binds to histone H1 protein and the promotors of histone H1 genes to regulate their expression. CRAMP1 is strongly localized to the histone locus body in the cell nucleus, however, CRAMP1 uniquely regulates only histone H1 expression. CRAMP1 interacts with another previously identified HLB component GON4L. The interaction between CRAMP1 and GON4L depends on the newly identified structured domains within CRAMP1, and the PAH repeats of GON4L.

The components of histone locus body and histone gene expression, including NPAT, GON4L and FLASH, are strongly conserved across species. The C-terminus of NPAT interacts with the SANT domains of both GON4L and FLASH^22,24^. Our analysis shows that GON4L also interacts with CRAMP1 via the PAH domains. Therefore, both GON4L and NPAT act as bridges between multiple components of the HLB complex. It is unclear if the NPAT, GON4L, CRAMP1 and FLASH exist as multiple distinct complexes or as a multisubunit complex. NPAT was observed in both co-IP and affinity purification of CRAMP1, suggesting that NPAT may be in the same complex as CRAMP1. Cell cycle timing of core histone gene expression is controlled by CyclinE/CDK2 of NPAT^21,26,52,53^. However, CRAMP1 is present at the promotors of the replication coupled histone H1 genes, H1-2 and H1-3, and the replication independent histone H1 gene H1-10. NPAT is not present at the H1-10 promoter but does accrue at low levels at the promoter of H1-2 and H1-4. NPAT plays a role in recruiting GON4L and FLASH to the HLB through their C-terminal SANT domains^22^. Phosphorylation of NPAT by Cdk2/CyclinE is required for transcriptional activation and maintenance of the histone genes at the G1/S boundary and during S-phase, respectively^18,21,52–54^.

There appears to be differential regulation of histone gene families. The HiNF-P/MIZF transcription factor binds to NPAT in its N-terminus and selectively regulates histone H4 expression^52,55^. The *Drosophila cramped* gene has been shown to localize to the histone locus body^56^. *Cramped* appears to be a selective regulator of histone H1, as mutations in *Drosophila* CRAMP led to severe reduction in the expression of histone H1, while having no effect on the core histones (H3, H4, H2A, H2B), implying a unique and conserved role for CRAMP1 in H1 regulation and/or stability.

The pattern of H1 variant gene expression differs by cell type and lineage. The components of the HLB, including CRAMP1 and NPAT, occupy the promoters of only the active genes that are in and open confirmation. Based on the correlation between increased DNA methylation and the closed chromatin state at histone genes, we propose that DNA methylation is the primary mechanism by which histone genes are selected for expression within a cell lineage.

## Acknowledgements

Thanks to the Northwestern Proteomics Core for analysis of the CRAMP1 affinity purifications. D.R.F was supported by R01GM143638 and U01CA260699. We thank Sui Huang and members of the Foltz lab for comments on the experiments in this manuscript. We thank Sudharsana Ravisankar for technical assistance in establishing the dTAG cell line.

## Materials and Methods

### Deletion mutants

A full length CRAMP1 entry clone was generated by BP Gateway recombination with C-terminal V5-tagged CRAMP1 Gateway expression clone (DNASU HsCD00865801) and pDONR221 (Invitrogen, 12536017) Gateway donor vector. A full length GON4L entry clone was synthesized de novo by Genewiz. CRAMP1 and GON4L deletion mutant entry clones were generated from their respective full length entry clones by site-directed mutagenesis. A C-terminal V5-tagged vector was generated using NEB HiFi Assembly by ligating a Gateway cassette into the large fragment produced from digesting full length CRAMP1 expression clone at BstBI and SpeI sites. Mutant CRAMP1 expression clones were generated by LR Gateway recombination reaction with mutant entry clones and a C-terminal V5-tagged vector. Mutant and full length GON4L expression clones were generated by LR Gateway recombination reaction with entry clones and a C-terminal FLAG-tagged vector. All plasmids were verified by sequencing and/or restriction digestion.

### Affinity Purification of CRAMP1

HEK293T stably over-expressing a CRAMP1-V5 fusion protein and parental HEK293T were lysed in extraction buffer (150 mM NaCl, 50 mM Tris-Cl pH 7.5, 1% v/v IGEPAL CA-630, protease inhibitor cocktail) for 30 minutes on ice. Whole cell extracts were centrifuged for 30 minutes at 4700 rpm at 4°C, supernatant collected and analyzed by BCA for protein concentration. The samples were normalized to the most dilute sample in the set, combined with magnetic V5-conjugated beads (Proteintech, v5tma), and incubated while rotating overnight at 4°C. Beads were washed 5x in wash buffer (150 mM NaCl, 50 mM Tris-Cl pH 7.5, 1% IGEPAL CA-630), after which beads were resuspended in 1X Laemmli buffer and boiled at 95°C for 10 minutes. Successful immunoprecipitation was tested by silver staining SDS-PAGE-resolved immunoprecipitated samples. Immunoprecipitated samples were trypsin digested and prepared for/subjected to mass spectrometry analysis. The Log2 fold change (FC) was calculated from normalized spectral counts of CRAMP1-V5 expressing cells versus parental. Three biological replicates of parental and CRAMP1-V5 -expressing cells were analyzed. P-values were calculated using t-test and corrected using Benjamini-Hochberg.

### Transient transfection and Co-immunoprecipitation (Co-IP)

HEK293T cells were cultured in Dulbecco’s Modified Eagle Medium (DMEM) supplemented with 10% heat-inactivated fetal bovine serum (FBS)-Optima (Bio-Techne, S12450) and 1% penicillin/streptomycin (Sigma, P0781). HEK293T plated in a 6 cm plate were transfected overnight in Opti-MEM with 1 µg DNA per construct (i.e. 2 µg DNA for co-transfection) using Lipofectamine 3000 reagent (Invitrogen, L3000015) and following the manufacturer’s protocol. Opti-MEM was replaced with growth media for 48 hrs. Cells were lysed in extraction buffer (150 mM NaCl, 50 mM Tris-Cl pH 7.5, 1% v/v IGEPAL CA-630, protease inhibitor cocktail) for 30 min on ice, centrifuged at 21,000 rcf for 30 min at 4°C. Only the supernatant was carried forward for Co-IP. Inputs were collected (5% total volume) and combined with sufficient 6X Laemmli buffer for a 1X dilution. Magnetic V5-conjugated (Proteintech, v5tma) or FLAG-conjugated beads (Sigma, M8823) were added to the remaining supernatant and incubated while rotating overnight at 4°C. Beads were washed 5x in wash buffer (150 mM NaCl, 50 mM Tris-Cl pH 7.5, 1% v/v IGEPAL CA-630) before suspension in 1X Laemmli buffer and boiling for 10 min at 95°C. Inputs and immunoprecipitated sample were analyzed by western blot. At least two unique biological replicates were analyzed per experiment.

### Structure prediction

The combined structure of GON4L and CRAMP1 complex was predicted as using Alphahold3 ^57^. Full-length sequences of both proteins were used for prediction although, only regions showing confidence pIDDT > 80 are shown in the structures. The interaction metric was derived from the expected position error by averaging the expected position error at each amino acid within or across molecules and reporting maximum expected position error minus the observed.

### Lentivirus production, transduction, and cell line generation

Lentivirus was generated in HEK293T by co-transfecting 12 µg desired transgene plasmid with 3 µg VSV-G coat protein vector, pMD2.G (Addgene, 12259), and 6 µg psPAX2 (Addgene, 12260) in a 10 cm plate. Parental cells were transfected overnight in Opti-MEM (Thermo Fisher, 31985070). Growth media was replaced, and the viral media was collected and replaced with fresh DMEM every 24 hr for two consecutive days. Viral media was pooled and sterile-filtered through a 0.45 µM surfactant-free cellulose acetate membrane filter. As needed, lentivirus was stored at −80°C or used immediately. Transduction was performed in 6-well format, where 5e5 parental cells were plated one day prior to infection. Lentivirus was spiked into growth media containing 8 µg/mL Polybrene (Sigma, H9268) and incubated overnight. Fresh growth media was replaced the next morning, and antibiotic selection was initiated 24 hr thereafter. Surviving colonies were pooled, and expression of the transgene was validated by immunoblotting for the exogenous tag.

### Immunoblot and Antibodies

Antibodies for HA (CST, 3724), V5 (CST, 13202), FLAG (Sigma, F1804), histone H1 (Thermo Fisher, PA5-30055), histone H3 (CST, 4459), ACTIN (Sigma, A2228), and GAPDH (Sigma, PLA0125) were purchased commercially. Samples for Western blot were prepared as indicated. For immunoprecipitated (IP) samples, the inputs and IP samples in extraction or wash buffer, respectively, were combined with 6X Laemmli buffer for a 1X working dilution, boiled at 95°C for 10 min, as described earlier. For non-IP samples, cells were lysed in radioimmunoprecipitation assay (RIPA) buffer (150 mM NaCl, 50 mM Tris-Cl pH 8.0, 1% v/v IGEPAL CA-630, 0.5% deoxycholate, 0.1% SDS) for 30 min on ice. Lysates were centrifuged at 20,000 rcf for 30 min at 4°C, and the supernatant was isolated and combined with sufficient 6X Laemmli buffer for a 1X working dilution, boiled at 95°C for 10 min. All samples were resolved by SDS-PAGE and transferred to polyvinylidene fluoride (PVDF) membrane, which was blocked in TBST supplemented with 10% non-fat milk powder for 20 min at room temperature. Primary antibody incubation was performed while rotating overnight at 4°C, and secondary antibody incubation was performed while rotating for 30 min at room temperature. One representative image of at least two replicated immunoblots is reported in the article.

### Biochemical fractionation and IP

HEK293T cells stably over-expressing CRAMP1-V5 were biochemically fractionated as described by Sebastien Gillotin, 2018. Magnetic V5 beads (Proteintech, v5tma) were added to nuclear and chromatin fractions. Immunoprecipitation was performed as described above, but a wash buffer identical to the particular extraction buffer was used. Samples were analyzed by Western blot.

### Real-time quantitative PCR

Total RNA was isolated using an RNA extraction kit (Qiagen, 74106) following the manufacturer’s protocol. Complementary DNA (cDNA) was prepared from 2 µg RNA using the iScript cDNA Synthesis kit (Bio-Rad, 17088900). cDNA was diluted 10-fold and combined with iTaq universal SYBR Green supermix (Bio-Rad, 1725121), then forward and reverse primers were added to a final concentration of 500 nM. The mixture was subjected to real-time quantitative PCR using the protocol described in the iTaq SYBR manual. Gene expression was analyzed by the 2^-ΔΔCt^ method. Three technical replicates were analyzed for every biological replicate, and a minimum of three biological replicates were analyzed for each experiment.

### ChIP-seq

ChIP-seq was performed in hTERT-RPE1 V5-CRAMP1 cells as previously described in Khurana *et al*, 2024. Briefly, 50×10^6^ cells were crosslinked with 1% formaldehyde, 1% FBS in 1XPBS for 10 min at room temperature with rotation, followed by quenching with 0.2M glycine for 5 min at room temperature with rotation. Cells were then washed in PBS, pelleted, and flash frozen in liquid nitrogen. Cells were lysed in lysis buffer 1 (50mM HEPES pH7.5, 140mM NaCl, 1mM EDTA, 10% glycerol, 0.5% IGEPAL CA-630, 0.25% Triton X-100), lysis buffer 2 (10mM Tris-HCl pH8.0, 20mM NaCl, 1mM EDTA, 0.5mM EGTA), and resuspended in lysis buffer 3 (10mM Tris-HCl pH8.0, 1mM EDTA, 0.1% SDS). Cells were sonicated using the Covaris E220 with the following parameters: 600 seconds, 10% duty cycle, 140 peak intensity and 200 cycles per burst. The lysates were centrifuged and input across the samples was normalized by DNA concentration. 10X ChIP dilution buffer (10% Triton X-100, 1M NaCl, 1% Na-Deoxycholate, 5% N-Lauroylsarcosine, 5mM EGTA) was added to a final concentration of 1X. 1% of the Input was removed and then DNA-protein crosslinked complexes were immunoprecipitated overnight at 4°C with the following antibodies: V5 (1.92ug CST), or Rabbit IgG (1.92ug CST). Complexes were then bound to Protein A Dynabeads (Invitrogen) for 4 hours at 4°C followed by 4 washes in RIPA buffer (50mM HEPES pH7.5, 500mM LiCl, 1mM EDTA, 1% IGEPAL CA-630, 0.7% Na-Deoxycholate) and 1 wash in 1X TE + 50mM NaCl. Reverse crosslinking reaction using proteinase K was performed at 65°C overnight, followed by DNA purification using Qiagen PCR purification kit. DNA libraries were produced using NEBNext Ultra II DNA Library Preparation kit and NEBNext Unique Dual Index Primers (New England Biolabs). Libraries were quality checked using Agilent D1000 TapeStation kit and Qubit Flourometer and then sequenced on the Illumina NovaSeq X (Admera Health) with 2×150 bp paired-end reads and with 1% PhiX as a control. We obtained 50M reads per sample. Data analysis including alignment of reads and peak calling was conducted using the nf-core/chipseq pipeline (https://github.com/nf-core/chipseq). Downstream analysis was performed using bedtools, deeptools and R. GO analysis of genes within 1kb of CRAMP1-consensus binding was conducted using Genomic Regions of Enrichment of Annotations Tools (GREAT) ^58^. NPAT ChIP-seq was conducted previously (GSE69147) ^51^.

### Modified reduced representation bisulfite sequencing

450,000 cells were lysed using QIAGEN RLT Plus buffer, and DNA isolation was performed with the QIAGEN AllPrep Micro Kit. Briefly, 200 ng of genomic DNA were subjected to restriction endonuclease digestion with MspI, and size selection, bisulfite conversion, and index ligation proceeded as previously described ^59–62^. The bisulfite conversion efficiency was 99.5% for all samples based on the observed methylation level of unmethylated λ-bacteriophage DNA (New England Biolabs) spiked in to each sample. Sequencing was performed on an Illumina NextSeq 2000 instrument with P2 kits (75-bp single-end reads). Processing, analysis, and quantification were performed using Trim Galore! v0.4.4, Bismark v0.16.3 with alignment to the hg38 genome build, and the Seqmonk bisulfite methylation pipeline as previously described ^63^. Visualization was performed by plotting raw methylation values from the Bismark bigwig file output and by generating cumulative distribution functions filtered to regions of open versus closed chromatin as defined by ATAC-seq analysis.

## Notes

### Competing Interest Statement

The authors have declared no competing interest.

